# Mathematical Model Predicts Tumor Control Patterns Induced by Fast and Slow CTL Killing Mechanisms

**DOI:** 10.1101/2023.07.19.548738

**Authors:** Yixuan Wang, Daniel Bergman, Erica Trujillo, Alexander T. Pearson, Randy F. Sweis, Trachette L. Jackson

## Abstract

Immunotherapy has dramatically transformed the cancer treatment landscape largely due to the efficacy of immune checkpoint inhibitors (ICIs). Although ICIs have shown promising results for many patients, the low response rates in many cancers highlight the ongoing challenges in cancer treatment. Cytotoxic T lymphocytes (CTLs) execute their cell-killing function via two distinct mechanisms: a fast-acting, perforin-mediated process and a slower, Fas ligand (FasL)-driven pathway. Evidence also suggests that the preferred killing mechanism of CTLs depends on the antigenicity of tumor cells. To determine the critical factors affecting responses to ICIs, we construct an ordinary differential equation model describing *in vivo* tumor-immune dynamics in the presence of active or blocked PD-1/PD-L1 immune checkpoint. Specifically, we identify important aspects of the tumor-immune landscape that affect tumor size and composition in the short and long term. By generating a virtual cohort with differential tumor and immune attributes, we also simulate the therapeutic outcomes of immune checkpoint blockade in a heterogenous population. In this way, we identify key tumor and immune characteristics that are associated with tumor elimination, dormancy, and escape. Our analysis sheds light on which fraction of a population potentially responds well to ICIs and ways to enhance therapeutic outcomes with combination therapy.

## 1 Introduction

Immunotherapy has launched a new era in cancer treatment that has remarkably improved outcomes for many patients. To have efficacy, immunotherapies must overcome immunosuppression induced by a tumor and its microenvironment, to allow the cytotoxic immune cells to target and kill cancer cells [4]. Immune checkpoint inhibitors (ICIs) are an important and well-studied class of immunotherapeutics that revitalize killing capacity of immune cells by blocking the activation of inhibitory immunoreceptors [13]. Immune checkpoint blockade therapy often results in a more durable response than chemotherapy or targeted therapies [13] and has shown promising results for many patients. However, the low overall response rates in many cancers present an ongoing challenge to clinicians. For example, objective response to checkpoint blockade monotherapy remains near 20% in patients with bladder cancer [21]. Over the past decade, there has been keen interest in research to improve the efficacy of blocking the programmed death-1/programmed death-ligand 1 (PD-1/PD-L1) immune checkpoint [29]. PD-1 is a protein expressed on a variety of immune cells and is highly expressed on tumor-specific T cells [11]. PD-L1 is expressed on some immune cells like T cells and B cells, and it is usually upregulated on tumor cells [18]. When PD-1 binds to PD-L1, the interaction inhibits the activation of tumor-infiltrating lymphocytes (TILs), induces the apoptosis of TILs, inhibits CTL killing activities, and promotes tumor metastasis and infiltration [18]. The FDA has approved seven monoclonal antibodies that target the PD-1/PDL-1 checkpoint with supplemental indications in over fifteen cancer types and two tissue-agnostic conditions [33]. Adding further complexity to the antitumor immune responses is the fact that CTLs execute their cell-killing function via at least two distinct mechanisms. The first process is fast-acting and perforin/granzyme mediated[12]. CTL-derived granzymes enter the tumor cell through perforin pores to induce structural damage and thus apoptosis [35]. The second process is a slower, FasL-driven killing mechanism [12]. FasL, a type II transmembrane protein upregulated on CTLs, can engage Fas on the target cell to trigger signaling that causes the activation of the apoptotic cascade [6]. In one study, perforin-based killing was detected within thirty minutes, whereas FasL-based killing was detected no sooner than two hours after tumor cell conjugated with CTL [12]. More recently, Cassioli and Baldari corroborated that granzyme/perforin mediated killing happen faster than Fas-dependent killing [6]. There is incomplete data regarding the manifestations of contact vs. granule-based T cell killing in solid tumors; however, some evidence suggests that CTLs switch from fast to slow killing with decreased antigen load [12], demonstrating different behaviors of the immune system towards tumor cells with varying antigenicity. The antigenicity of a tumor can be defined as the extent to which tumor cells display HLA-restricted antigens that can be selectively or specifically recognized by immune cells [28]. One of the immunoevasive strategies cancer cells employ is the downregulation or loss of antigens, which can occur due to the immune selection of cancer cells that lack immunogenic tumor antigens and through the acquisition of defects in antigen presentation [2]. Loss of antigenicity impairs the ability of natural immune responses to control cancers, impedes immunotherapies that work by re-invigorating CTLs, and potentially alters future responsiveness to additional treatments [9].

By constructing and analyzing data-driven mathematical and computational models, we aim to investigate the roles of antigenicity, differential immune cell-kill mechanisms and other aspects of the tumor-immune landscape in immune checkpoint blockade therapy, and to formulate therapeutic strategies to enhance outcomes. The complex interactions between tumor and the immune system result in vastly different outcomes such as tumor elimination, tumor dormancy and uncontrolled tumor growth. One way to predict the specific circumstances leading to these fates is through non-linear ordinary differential equations (ODEs), which model cellular interactions and reveal the temporal dynamics of the components in tumor-immune interactions. In what has become a classic model of tumor immune dynamics, Kuzentzov proposed a system of ODEs that model the cytotoxic T lymphocyte response to the growth of an immunogenic tumor[19]. Key features of this model include the characterization of antigen-independent and antigen-stimulated recruitment of cytotoxic effector cells to the tumor site, and an immune-mediated death rate of tumor cells that increases proportionally as the number of T cells increases. Alternative mathematical descriptions of tumor cell kill, such as the “Beddington” functional response [14], have now been proposed to reflect the more realistic assumption that immune cell killing saturates as the number of immune cells increases.

In this study, we test the hypothesis that immune checkpoint therapy impacts both the total tumor volume and the proportion of the two tumor cell phenotypes due to differential killing mechanisms associated with each tumor phenotype. The full details of the model formulation are in Section 2, followed by sensitivity analysis, bifurcation analysis and virtual cohort simulations in Section 3. Our analysis highlights important parameters that affect the outcomes of immune checkpoint blockade. Among these, some parameters characterize the tumor-immune landscape, and others can be modulated by cancer treatments such as chemotherapy and additional types of immunotherapy. Understanding the impact of different parameters on tumor volume and phenotypic composition enables us to gain insights into which patients are most likely to respond to ICIs and what combination therapy can result in better therapeutic outcomes.

## 2 Model Formulation

Our mathematical model reflects the current biological understanding of tumor-immune interactions including the role of the PD-1/PD-L1 immune checkpoint. We develop a “checkpoint active” model that describes the fully functional PD-1/PD-L1 signaling pathway leading to immune suppression. We then relax this assumption by removing the immunosuppressive effects by the PD-1/PD-L1 complex to formulate a model with the PD-1/PD-L1 signal inhibited to study the implications of a perfect checkpoint blockade therapy. The model captures the temporal evolution of the number of high antigen tumor cells (*N*), low antigen tumor cells (*M*) and cytotoxic T cells (*T*). Model variables and their units are described in Table 1.

**Table 1:** Variables for the ODE model with active immune checkpoint. We use the conversion factor 10^6^ tumor cells equal 1mm^3^ [8].

| **Variable** | **Description** | **Units** |
| --- | --- | --- |
| $N$ | Tumor cells with high antigencity | # |
| $M$ | Tumor cells with low antigencity | # |
| $T$ | Cytotoxic T cells | # |

Figure 1 provides a schematic diagram of the components of the ODE model.

**Figure 1:**
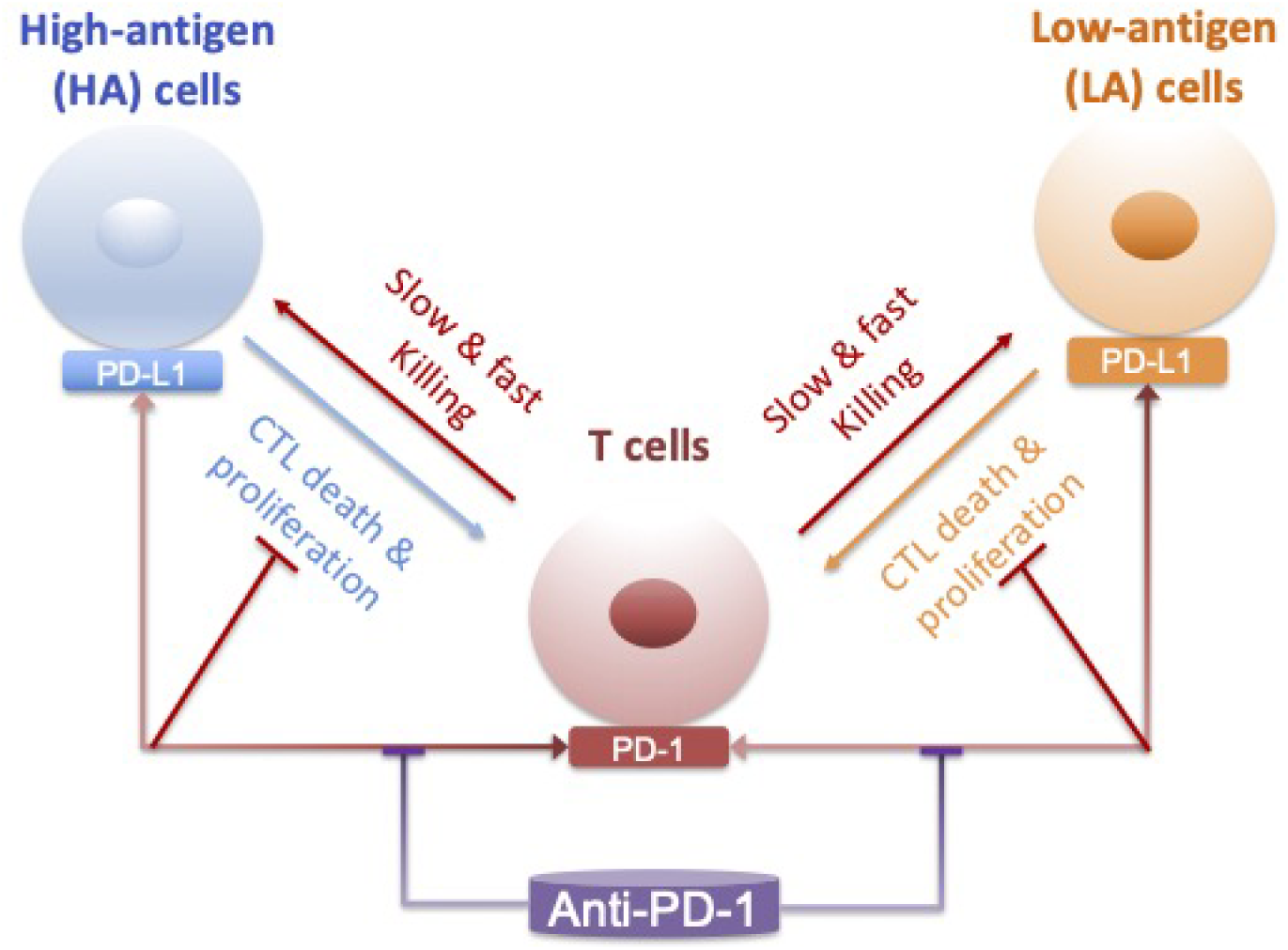
Schematic diagram of the ODE model describing tumor-immune dynamics. The two tumor cell phenotypes are high antigen (HA) cells and low antigen (LA) cells. T cells kill tumor cells via two mechanisms: a fast-acting and perforin/granzyme-mediated process, and a slower, Fas ligand (FasL)-driven process. PD-1 expressed on T cells and PD-L1 expressed on tumor cells interact to inhibit T cell activity in tumor killing. Anti-PD-1 prevents the engagement of PD-1 and PD-L1.

We consider a heterogeneous tumor made up of tumor cells with both high and low antigenicity. Equation (1) models the temporal dynamics of the tumor cells that express high antigen levels.

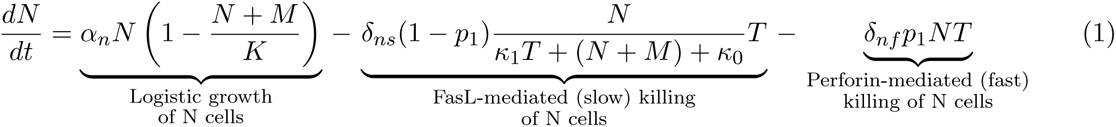

The first term in Equation (1) describes tumor cells with high antigenicity proliferating logistically with growth rate *α_n_* and carrying capacity *K*. This logistic form reflects the implicit competition for space and resources that occurs between the two cell types. The second term in Equation (1) describes the slow killing of high antigen tumor cells with probability 1 *− p*_1_ and maximum rate *δ_ns_*. Here, slow killing is modeled by the “Beddington” function similar to the saturating function used in [15]. We assume that each activated CTL kills multiple tumor cells, independent of their antigenicity, primarily by inducing apoptosis in a contact dependent manner. The third term in equation (1) describes the faster, perforin-mediated killing of high antigen tumor cells with probability *p*_1_ and maximum rate *δ_nf_*. We model rapid cell killing as being directly proportional to the numbers of CTLs and tumor cells as described in [19]. There is some evidence that, with strong antigenic stimulation, there can be a substantial proportion of high-rate killer CTLs [34]. Although this could imply that high antigen tumor cells are preferentially killed via the fast mechanism, we keep the model general and assume that high antigen tumor cells can be killed via both the fast perforin-mediated pathway and the slow FasL-mediated pathway in order to test the differential impact of cell kill decisions.

Equation (2) models the temporal dynamics of the tumor cells that express low antigen levels.

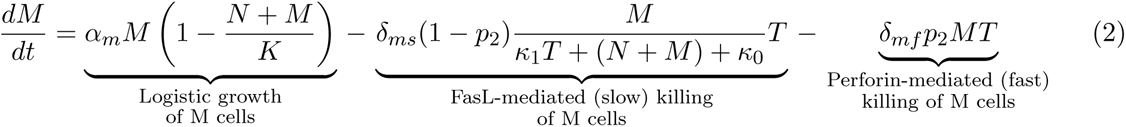

Although quantitative insights into the dynamic cell kill patterns of CTLs are essential for the rational design of immune-based therapies, the CTL killing landscape with a growing tumor remains insufficiently characterized [3]. There is evidence that sustained killing at the population level, relies on a highly variable, multiple-killing performance at the individual CTL level [34]. Therefore, we decided to model the growth and killing of low antigen tumor cells just like the high antigen tumor cells, with necessary changes to the names of the parameters. By allowing maximum flexibility in which type of tumor cells are predominantly killed by each mechanism, we are able to investigate the impact of probabilities for fast (*p*_1_*, p*_2_) and slow kill (1 *− p*_1_, 1 *− p*_2_) on the final tumor volume and composition. However, we anticipate, based on [34], that *p*_1_ *> p*_2_, i.e. high antigen tumor cells are more likely to be killed via the fast mechanism than low antigen tumor cells.

In our model the CTLs we consider are activated CD8+ cytotoxic T cells, which are terminally differentiated effector T cells with cytotoxic activity. Equation (3) models the temporal dynamics of these activated CD8+ T cells.

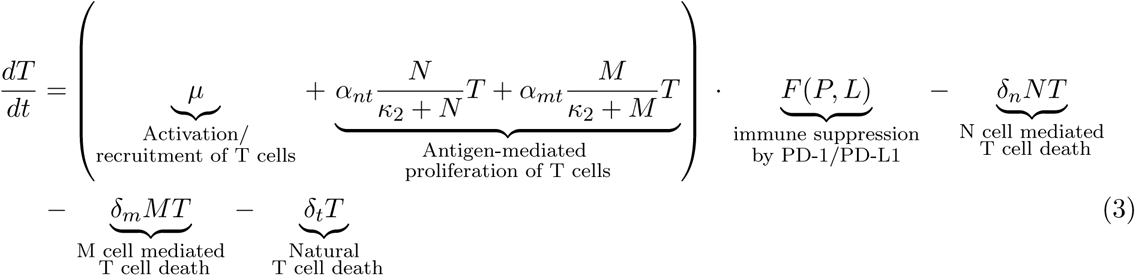

The first term in (3) represents a constant recruitment/activation of T cells at rate *µ*. The second and third terms describes proliferation that occurs as the result of antigenic stimulation by the tumor cells. We assume both tumor phenotypes can elicit proliferation of T cells, although we anticipate *α_nt_ > α_mt_*. *F* (*P, L*) models the immune suppression by the PD-1/PD-L1 complex. As in [20, 22, 31, 23], the function for suppression of T cell activation and proliferation by the PD-1/PD-L1 complex is given by:

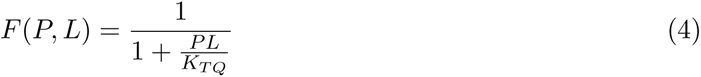

where *P* and *L* represent the concentrations of PD-1 and PD-L1 respectively. This formulation ensures that as PD-1 and PD-L1 increase, so does the number of PD-1/PD-L1 complexes. An increase in these immune checkpoint complexes corresponds to a smaller value of *F* (*P, L*), thereby modeling the inhibition of T cell activity. Finally the last three terms in equation (3) describe the ways in which CTLs die. Specifically, interaction with high antigen expressing cells can result in death at rate *δ_n_*, interaction with low antigen expressing cells can result in death at rate *δ_m_*, and CTLs can die naturally at rate *δ_t_*.

The checkpoint active model described above serves as the baseline case, which can be modulated by adjusting the immunosuppressive effects. We will consider the special case of 100% effective checkpoint blockade therapy, which corresponds to *F* (*P, L*) = 1 because it can provide insights about the best-case scenario outcomes for therapies targeting PD-1 or PD-L1. Baseline parameters in Table 2 are chosen from literature where possible. Due to the novel components in our model, such as different cell kill mechanisms for high and low antigen tumor phenotypes, we chose parameters not available in the literature so that the total tumor volume on Day 25 after complete checkpoint blockade is reduced by 75%, compared to the checkpoint active case.

**Table 2:** Baseline parameters.

| Parameter | Description | Value (Baseline) | Units | Source |
| --- | --- | --- | --- | --- |
| $\alpha_n$ | Proliferation rate of high antigen tumor cells | 0.05 - 0.8 (0.337) | per day | Estimated ([23]) |
| $\alpha_m$ | Proliferation rate of low antigen tumor cells | 0.05 - 0.6 (0.337) | per day | [23] |
| K | Carrying capacity for tumor cells | $3 - 6 \cdot 10^9 (5 \cdot 10^9)$ | # of cells | Estimated |
| $\delta_{ns}$ | Maximum CTL-induced death rate of high antigen tumor cells via the slow killing mechanism | 1 - 12 (4) | per day | Estimated ([5]) |
| $\delta_{ms}$ | Maximum CTL-induced death rate of low antigen tumor cells via the slow killing mechanism | 1 - 12 (4) | per day | Estimated ([5]) |
| $\delta_{nf}$ | CTL-induced death rate of high antigen tumor cells via the fast-killing mechanism | $10^{-8} - 10^{-6} (2.5 \cdot 10^{-7})$ | per cell per day | Estimated ([23]) |
| $\delta_{mf}$ | CTL-induced death rate of low antigen tumor cells via the fast-killing mechanism | $10^{-8} - 10^{-6} (2.5 \cdot 10^{-7})$ | per cell per day | Estimated ([23]) |
| $\kappa_0$ | Half-saturation constant in maximum CTL-induced death rate of via the slow killing mechanism | $10^6 - 10^8 (2 \cdot 10^7)$ | # of cells | Estimated |
| $\kappa_1$ | dimensionless | 0.01 - 1 (0.5) | dimensionless | Estimated ([19]) |
| $\kappa_2$ | Half-saturation constant in the antigen-mediated T cell proliferation rate | $10^6 - 10^8 (2.019 \cdot 10^7)$ | # of cells | Estimated ([23]) |
| $\delta_t$ | Death rate of T cells | 0 - 0.5 (0.0412) | per day | [22] ([19]) |
| $\mu$ | Activation and recruitment rate of T cells | $5 \cdot 10^3 - 1.5 \cdot 10^5 (2 \cdot 10^4)$ | # per day | Estimated |
| $\delta_n$ | CTL death rate due to interactions with high antigen tumor cells | $10^{-11} - 10^{-9} (3.422 \cdot 10^{-10})$ | per cell per day | Estimated ([19]) |
| $\delta_m$ | CTL death rate due to interactions with low antigen tumor cells | $10^{-11} - 10^{-9} (3.422 \cdot 10^{-10})$ | per cell per day | Estimated ([19]) |
| $p_1$ | Probability of high antigen tumor cells death via the fast-killing mechanism | 0 - 1 (0.92) | dimensionless | Estimated |
| $p_2$ | Probability of low antigen tumor cells death via the fast-killing mechanism | 0 - 1 (0.33) | dimensionless | Estimated |
| $\alpha_{nt}$ | Maximum rate of CTL proliferation activated by N cells | 0 - 0.5 (0.15) | per day | Estimated |
| $\alpha_{mt}$ | Maximum rate of CTL proliferation activated by M cells | 0 - 0.5 (0.15) | per day | Estimated |
| $\rho_p$ | Molar concentration of PD-1 per T cell | $10^{-12} - 10^{-10} (1.259 \cdot 10^{-11})$ | $\mu M$ | [31], [22] |
| $\rho_l$ | Molar concentration of PD-L1 per T cell | $10^{-12} - 2 \cdot 10^{-10} (2.510 \cdot 10^{-11})$ | $\mu M$ | [31], [22] |
| $\epsilon_c$ | Expression of PD-L1 on tumor cells vs. T cells | 1 - 50 (10) | dimensionless | [31] |
| $\mu_{PA}$ | Blocking rate of PD-1 by anti-PD-1 | 6.45 - 273 (8.945) | L/$\mu$ mol/h | [31], [20] |
| $k_{TQ}$ | Inhibition of T cells by PD-1/PD-L1 | $10^{-10} - 10^{-8} (1.296 \cdot 10^{-9})$ | $(\mu M)^2$ | [31], [20] |

## 3 Results

We use mathematical and computational tools, including sensitivity analysis, bifurcation analysis, and virtual cohort simulations, to understand how model parameters affect tumor growth and response to checkpoint blockade therapy or combination therapy.

### 3.1 Sensitivity of tumor growth to parameters characterizing the tumor-immune landscape

Many details of tumor-immune interactions in the presence or absence of the PD-1/PD-L1 immune checkpoint remain unknown or hard to quantify, leading to uncertainty in model parameters and initial conditions. We use global sensitivity analysis to understand the impact of this uncertainty and to determine which parameters have the greatest impact on tumor growth when the PD-1/PDL1 immune checkpoint is active and when it is completely blocked. Following [23], we assess global sensitivity by using Latin hypercube sampling (LHS) and the partial rank correlation coefficient (PRCC) between the parameters and the tumor volume on Day 25 and Day 150. Sensitive parameters are defined to be those in the upper quartile of all PRCC values in terms of magnitude and with a p-value of less than 0.05. Figure 2A shows the PRCC between the parameters and the tumor volume when the checkpoint is active. The tumor volume is most sensitive to *α_n_, α_m_*, *K*. On Day 25, most tumors are still growing and have not reached the carrying capacity. Therefore, tumor proliferation rates (*α_n_, α_m_*) have higher PRCC value on Day 25 than Day 150. With checkpoint active, most tumors have reached the carrying capacity on Day 150, which explains the nearly perfect positive correlation between tumor volume and the carrying capacity (*K*).

**Figure 2:**
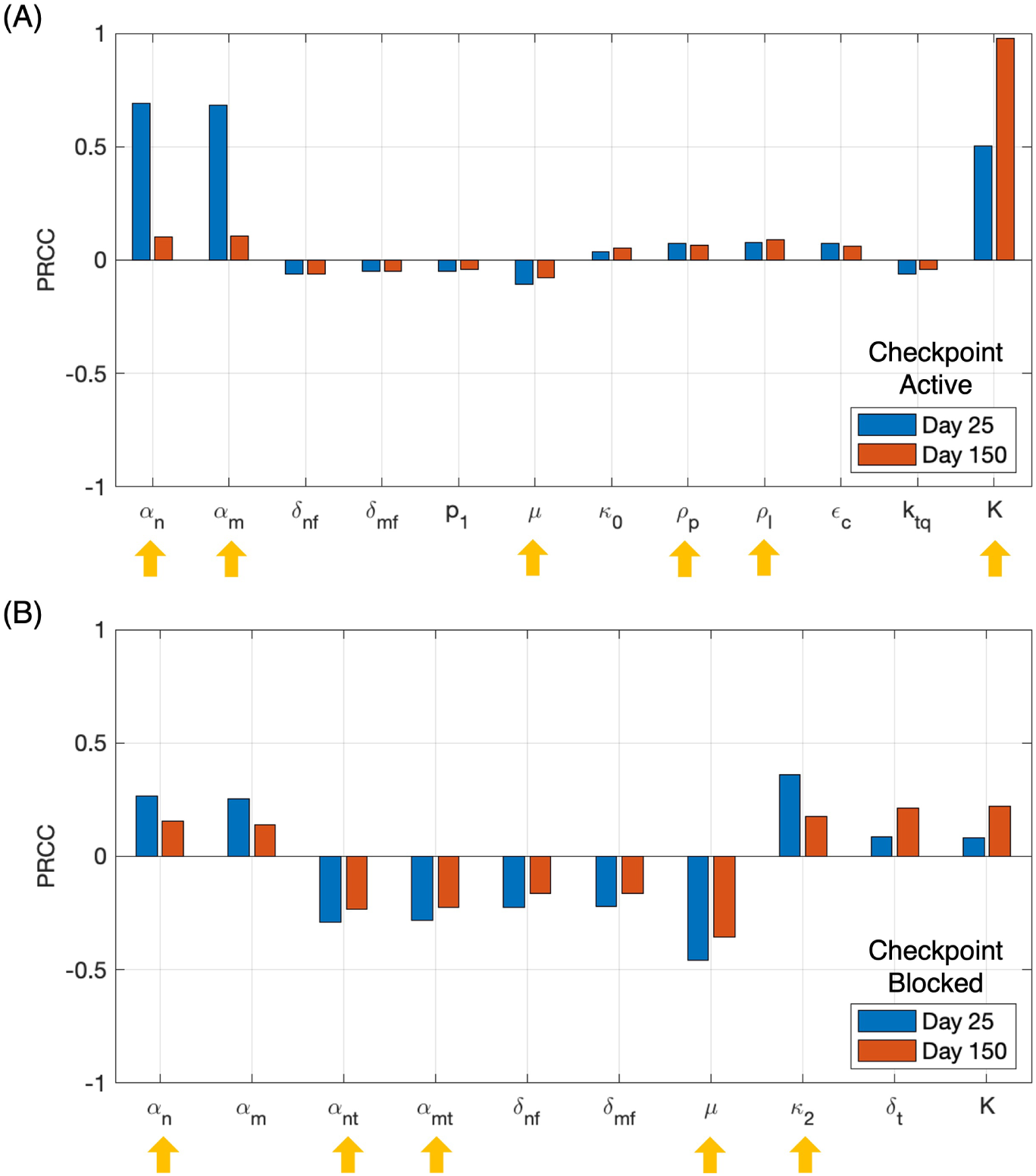
Immune checkpoint activity affects the most sensitive parameters with respect to tumor volume. (A) PRCCs (partial rank correlation coefficient) of parameters in the model with immune checkpoint active. (B) PRCCs of parameters in the model with immune checkpoint blocked. Blue: PRCC with respect to total tumor volume on Day 25. Red: PRCC with respect to total tumor volume on Day 150. Yellow arrows: sensitive parameters with magnitude of PRCC ranked in the top quartile. Parameters with magnitude of PRCC ranked in the top 50% are shown.

When the PD-1/PD-L1 checkpoint is blocked, there is a different set of sensitive parameters as the PRCC results in Figure 2B show. The total tumor volume is still sensitive to *α_n_, α_m_* (tumor proliferation rates) but less sensitive to *K* (carrying capacity). When the checkpoint is blocked, the total tumor volumes are in general smaller and well below the carrying capacity, resulting in the smaller PRCC value for *K*. The tumor volume is now most sensitive to *µ* (activation and recruitment rate of T cells), followed by *κ*_2_ (half-saturation constant in antigen mediated T cell proliferation), *α_nt_, α_mt_* (maximum rate of antigen-mediated CTL proliferation), *α_n_, α_m_*, *δ_nf_, δ_mf_*(CTL-induced death rate via fast killing), *K* and *δ_t_* (death rate of T cells). The total tumor volume is negatively correlated with *α_nt_, α_mt_, δ_nf_, δ_mf_, µ* and positively correlated with *κ*_2_*, α_n_, α_m_, K, δ_t_*. The results show that any therapy that enhances antigen-independent T cell recruitment, increases the rates of either type of killing mechanism, or increases antigen-mediated T cell proliferation may be combined with immune checkpoint blockade therapy to effectively reduce tumor size.

We conducted similar analysis (Figure S1) to determine which parameters have the greatest impact on tumor composition, which we measure by the ratio of low antigen tumor cells to total tumor cells. The endpoints of PRCC analysis is now low antigen to total tumor cell ratio on Day 25 and Day 150. In this case, with checkpoint blocked or active, ratio of low antigen to total tumor cell is sensitive to the same set of variables: *α_n_, α_m_, δ_nf_, δ_ns_, p*_1_ (probability of high antigen cell death via fast killing), *p*_2_ (probability of low antigen cell death via fast killing) and *R_ha_*(initial ratio of high antigen tumor cells to total tumor cells). In particular, *p*_1_ and *p*_2_ are new sensitive parameters that are important mediators of the ratio of low antigen tumor cells to total tumor cells.

### 3.2 Impact of immune responses parameters on long-term tumor volume and composition after complete checkpoint blockade

Many parameters of our model are crucial because they either characterize the distinct immune cell-kill mechanisms or can be modulated with therapy. By changing some of these parameters in the model, we now study the impact of the different cell-kill mechanisms (*p*_1_*, p*_2_) and individual or combined therapies. Examples of treatment strategies include immune checkpoint inhibition (*F* (*P, L*) = 1), stimulated immune cell expansion due to the administration of therapeutic cytokines like IL-2 (increases in *α_nt_, α_mt_*) [17], and adoptive T-cell transfer (increasing *µ*). Moreover, the fact that, according to Section 3.1, tumor volume and composition are sensitive to these parameters further necessitates analysis of model predictions when these parameters vary.

Bifurcation analysis studies the changes in the number and stability of steady states as model parameters change. We use bifurcation analysis to investigate how stable steady state tumor volume and composition change in response to variations in sensitive parameters when the PD-1/PD-L1 checkpoint is completely blocked. We plot two and three-parameter bifurcation diagrams to reveal important relationships between parameters associated with T cell killing mechanism (*p*_1_*, p*_2_) and those that can be modulated by treatment (*µ, α_nt_, α_mt_*).

#### Differential immune cell-kill mechanisms determine long-term tumor composition

Figure 3A is a two-parameter bifurcation diagram that shows how tumor composition in a tumor-persistent steady state varies depending on the values of *p*_1_ and *p*_2_. When a tumor of any size persists at steady state, the *p*_1_ *− p*_2_ parameter space is dissected into regions where the tumor either consists of only high antigen tumor cells or only low antigen tumor cells, represented by striped and solid regions, respectively. These high antigen-dominant and low antigen-dominant scenarios are the only two types of nonzero steady state outcomes for tumor composition predicted by our model for the parameter ranges in Table 1. Figure 3A shows that when the probability of high antigen tumor cell death via the fast-killing mechanism is greater than that for low antigen tumor cells (*p*_1_ *> p*_2_), the final tumor composition at steady steady will be homogeneous and only consists of low antigen cell; and vice versa.

**Figure 3:**
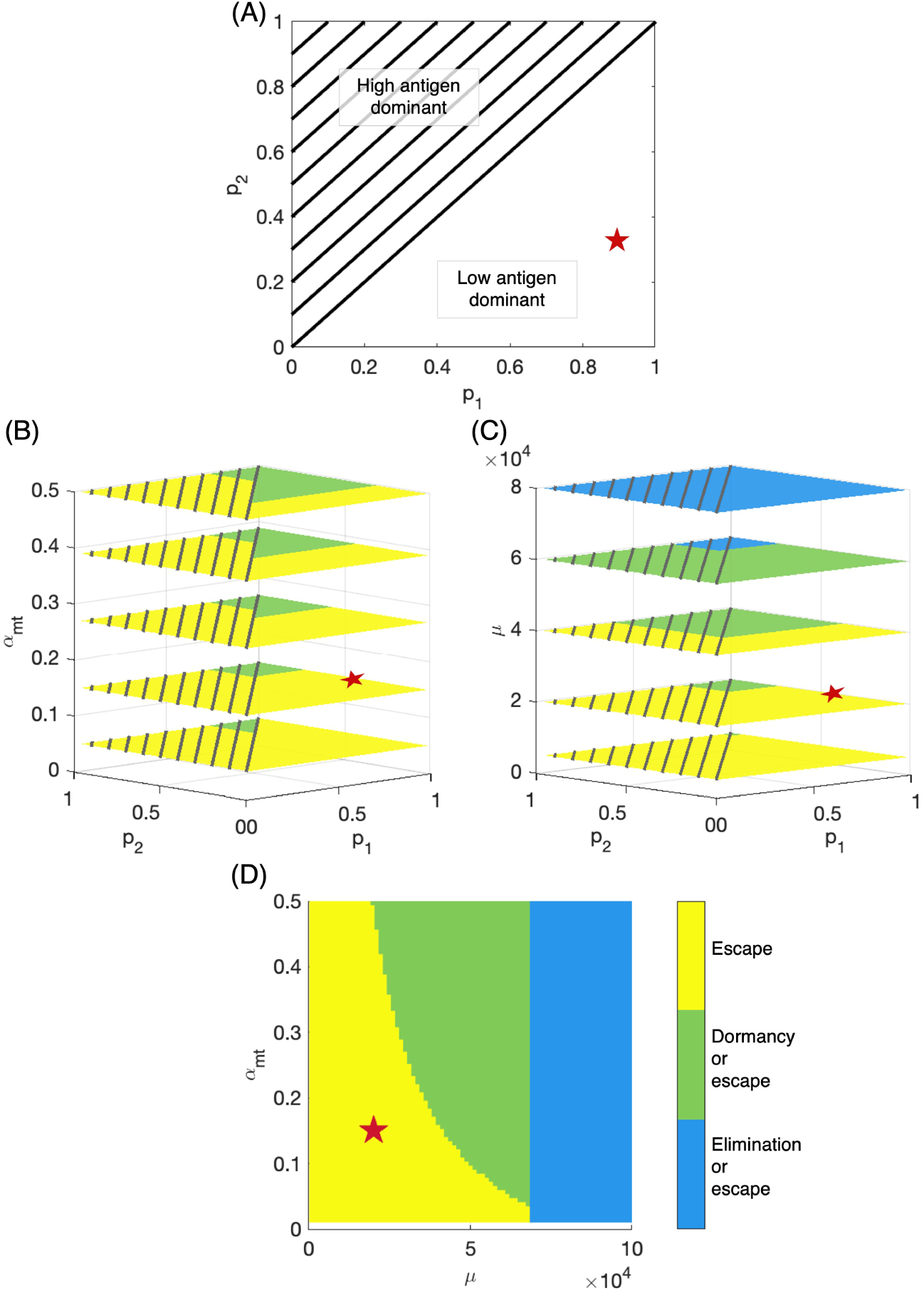
Bifurcation analysis illustrates important variations in steady state tumor size, composition, and response to ICI monotherapy or combination therapy. (A) Steady state tumor composition in relation to *p*_1_*, p*_2_ values. High (low) antigen dominant: consisting of only high (low) antigen tumor cells. (B) Response to changing *α_mt_* (e.g. cytokine therapy) at various *p*_1_*, p*_2_ values. Two outcomes in terms of tumor size: escape (yellow) and bistability with dormancy or escape (green).(C) Response to changing *µ* (e.g. adoptive T cell transfer) at various *p*_1_*, p*_2_ values. One more outcome in addition to panel B: bistability with elimination or escape (blue). (D) Comparison of the impact of *µ* and *α_mt_* on steady state tumor size. Escape: tumor size *≥* 500mm^3^; dormancy: 0 *<* tumor size *<* 500mm^3^; elimination: tumor size = 0mm^3^. Red star: baseline parameters. Parameters not shown are fixed at baseline values.

#### Combination checkpoint blockade and cytokine therapy enhances opportunities for tumor dormancy

Our model predicts three types of outcomes in terms of steady state tumor size: elimination, dormancy, and escape. Elimination is defined as no tumor cells present in the system. Dormancy is characterized by the tumor approaching a small, nonzero steady state, and escape is characterized by the tumor approaching carrying capacity. In addition, our model also predicts bistability, which occurs when the steady state tumor can be any two among the three outcomes (e.g. elimination and dormancy) depending on the initial conditions.

Fixing all other parameters at baseline values, we explored the impact of *p*_1_*, p*_2_, and *α_mt_* on the realization of these outcomes. Figure 3B considers the *p*_1_*−p*_2_ bifurcation plane for five values of *α_mt_*, the maximum rate of CTL proliferation activated by low antigen tumor cells. The slices are taken at *α_mt_* = 0.05, 0.15, 0.27, 0.39, 0.5, with the second from the bottom slice representing our baseline *α_mt_*. This figure shows that there are two types of therapeutic outcomes in terms of steady state tumor size: escape and bistability, represented by yellow and green, regions, respectively. The bistability region in this case includes both dormancy and escape as possible outcomes. With baseline parameters (red star in Figure 3B) and the PD-1/PD-L1 checkpoint completely blocked, the tumor will escape and grow to carrying capacity. However, an additional therapy (e.g. cytokine therapy) that increases *α_mt_* makes tumor dormancy possible. More generally, our model predicts that a therapy that modulates *α_mt_* shrinks the region of *p*_1_ *− p*_2_ values where the tumor escapes with certainty and expands the region where the tumor can potentially stabilize in a small, dormant state. It is noted that steady state tumors in Figure 3B will be either very large (escape) or small (dormant) and either completely high (*p*_1_ *< p*_2_) or low (*p*_1_ *> p*_2_) antigen dominant. One would observe the same trend if *α_nt_* is used instead of *α_mt_*because high and low antigen tumor cells are modelled in the same way mathematically.

#### Combination checkpoint blockade and adoptive T-cell therapy enhances opportunities for tumor elimination

Similarly, Figure 3C considers the *p*_1_*−p*_2_ bifurcation plane for five values of *µ*, the T cell recruitment and activation rate. The slices are taken at *µ* = (0.5, 2, 4, 6, 8) *×* 10^4^, with the second from the bottom slice representing our baseline *µ*. This figure again shows that the yellow region, representing escape, shrinks as *µ* increases. However, as *µ* increases further, a third outcome emerges, bistability resulting in elimination or escape represented by the blue regions in Figure 3C. These results imply that an additional therapy that increases *µ* (e.g. adoptive T-cell therapy) can potentially, result in tumor clearance. With baseline parameter values (red star in Figure 3C) and the PD-1/PD-L1 checkpoint completely blocked, the tumor will grow to carrying capacity. In this case, increasing *µ* can result in tumor dormancy or even elimination. Overall, increasing *µ* shrinks the region of *p*_1_ *− p*_2_ values where the tumor escapes with certainty and expands the region where the tumor has a chance of being eliminated or stabilizing in a small, dormant state.

#### Pre-treatment immune landscape determines the feasibility of combination therapeutic strategies

Taken together, the results lead us to investigate the potential of combining cytokine and adoptive T cell therapies by exploring the *µ−α_mt_* plane. Figure 3D shows how this type of combination therapy impacts therapeutic outcomes when *p*_1_*, p*_2_ are fixed at their baseline values. Here we see that if a tumor is characterized by small intrinsic values of *µ* and *α_mt_* that would normally lead to escape (the lower left corner of the graph), the fastest way to move to a potentially better outcome with the least amount of additional treatments is combination therapy (checkpoint blockade, adoptive T-cell transfer and cytokine therapy) that simultaneously increases both *µ* and *α_mt_*. Figure 3D also implies that it is now possible with combination therapy (checkpoint blockade and adoptive T-cell therapy) to only increase *µ* and move from any location in the yellow escape region to the blue region where elimination is possible. If a tumor is characterized by our baseline parameter values represented by the red star in Figure 3D, then additional cytokine therapy within the range for *α_mt_* that we consider will be ineffective for changing the therapeutic outcome of escape, but if *α_mt_* can be increased to approximately four times our baseline value, dormancy is possible.

### 3.3 Clinical implications of variable probabilities of fast killing

The previous subsection examined how steady state outcomes for tumor size vary with parameters that can be modulated by therapy as the probability of tumor cell death via fast killing changes. These steady states are rarely seen in practice in the pre-clinical setting. To consider these effects on clinically relevant timeline, we measure the total tumor volume on Day 25 and Day 150. The first end point of 25 days was chosen because model tumor xenograft experiments in mice are often conducted for 3-4 weeks. The second endpoint of 150 days was chosen to represent the timescale of clinical elimination. Figure 4 shows the changes in total tumor volume and composition on Day 25 and Day 150 for varying *p*_1_*, p*_2_ combinations when the PD-1/PD-L1 checkpoint is active or completely blocked. Again, *p*_1_ is the probability of high antigen tumor cell death via the fast-killing mechanism and *p*_2_ is the probability of low antigen tumor cell death via the fast-killing mechanism. Due to limitations of current imaging technology, tumors smaller than 0.1mm^3^ cannot be detected [26]. Therefore, we define clinical elimination as a tumor smaller than 0.1mm^3^. For the same reason, we stop our simulations when the total tumor size reaches 0.1mm^3^ or below. When this happens, the tumor size remains at 0.1mm^3^. The size of a circle in Figure 4 represents the size of the tumor and the color of a circle is a measure of tumor composition as it represents the ratio of low antigen tumor cells to total tumor cells. As the tumors move up the color bar from blue to red, they transition from high antigen-dominant to low antigen-dominant.

**Figure 4:**
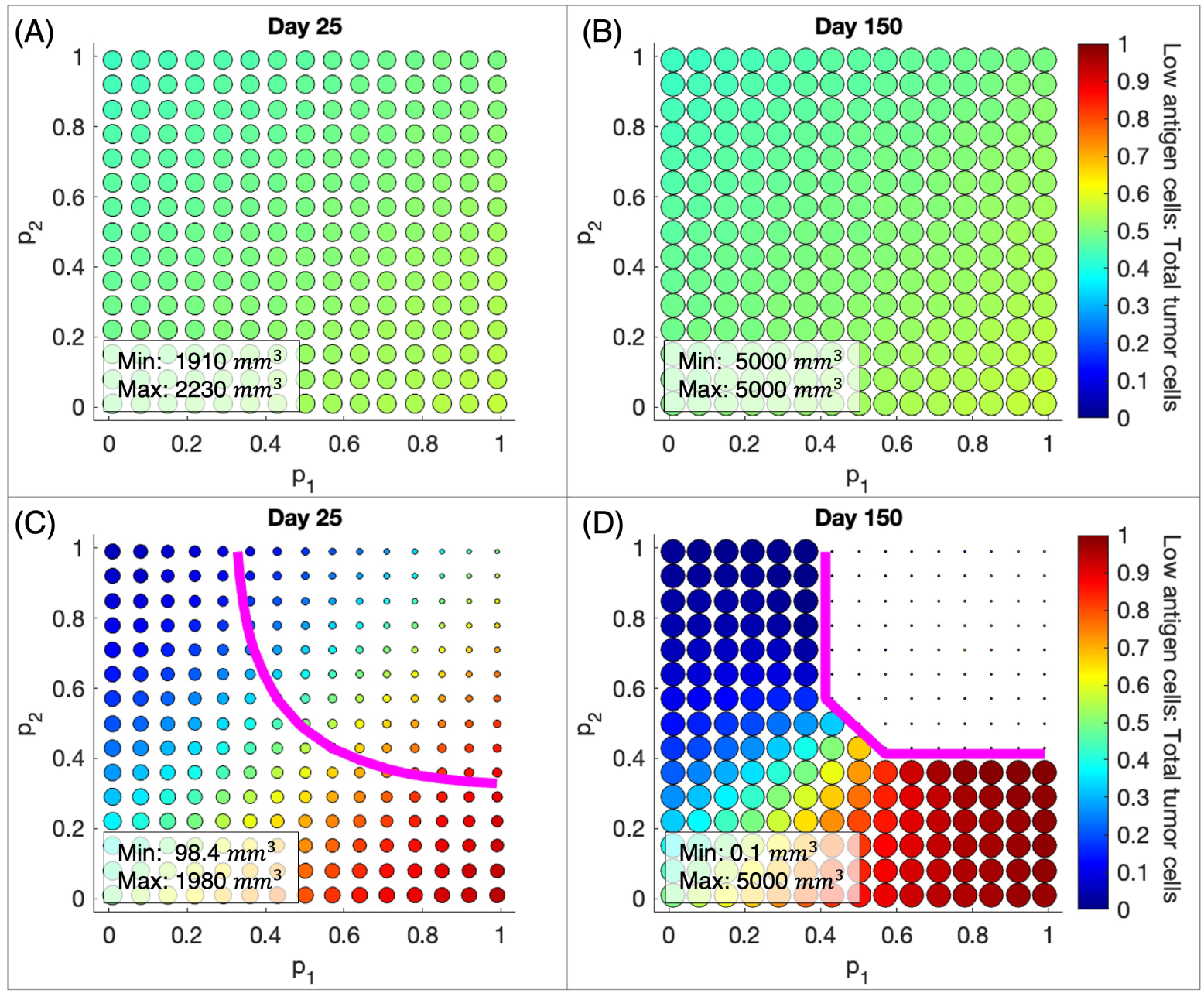
Probability of “fast” T cell killing determines the dominant cell type and tumor size in response to checkpoint blockade therapy. (A), (B) Tumor size and composition with checkpoint active on Day 25 and Day 150. (C), (D) Tumor size and composition with checkpoint blocked on Day 25 and Day 150. Magenta line: combinations of *p*_1_*, p*_2_ that result in 75% reduction in total tumor volume after checkpoint blockade, relative to when the checkpoint is active. Size of the circles: tumor size; color of circles: the ratio of low antigen tumor cells to total tumor cells, a measure of tumor composition. Initial tumor size in all simulations is 1 mm^3^ and consists of 50% high antigen tumor cells and 50% low antigen tumor cells.

Figure 4A shows that as *p*_1_*, p*_2_ vary when the checkpoint is active, the total tumor volume on Day 25 remains largely unaffected; all tumors are within about 300mm^3^ of each other. By Day 150, these tumors have all reached the maximum carrying capacity volume (Figure 4B). Tumor composition varies little when the checkpoint is active despite large variations in *p*_1_*, p*_2_. The initial tumor size is 1mm^3^ and consists of 50% high antigen tumor cells and 50% low antigen tumor cells. As the difference between *p*_1_ and *p*_2_ increases, moving towards the lower right corner, there is a modest shift toward low antigen dominance with low antigen tumor cells making up at most 56% of the tumor cells. Similarly, moving towards the upper left corner, there is a modest shift toward high antigen dominance.

Outcomes for complete checkpoint blockade are shown in Figures 4C and 4D. When *p*_1_ *> p*_2_, the proportion of low antigen tumor cells in the tumor increases significantly after checkpoint blockade, as illustrated by the shift to more red colors in the lower right portion of the graphs. The magenta curve in Figures 4C and 4D represents the combinations of *p*_1_*, p*_2_ that result in 75% reduction in total tumor volume and our baseline values fall into this region. These figures demonstrate the importance of the probability of fast killing on the therapeutic outcomes of PD-1/PD-L1 checkpoint blockade. If both cell types are killed by the fast mechanism, the resulting tumor on Day 25 is twenty times smaller than if both cell types are killed by the slow mechanism, as illustrated by the difference in size of the dots in the upper right and lower left corners. If high antigen tumor cells are killed only via the fast mechanism, then low antigen tumor cells must have at least 0.33 probability of being killed via the fast mechanism in order to achieve at least 75% tumor reduction on Day 25. The relative values of *p*_1_*, p*_2_ determine the prevailing type of tumor cells in the long run. When *p*_1_ *> p*_2_, checkpoint blockade therapy will increase the proportion of low antigen phenotype in the tumor despite reducing the total tumor size. This change in tumor composition will impact the tumor’s responsiveness to future treatments. For example, since low antigen tumor cells in our baseline assumption elicit slower immune response, a more low antigen-dominant tumor will likely be less responsive to subsequent immunotherapy.

Figure 4D shows that by Day 150 several *p*_1_*, p*_2_ combinations lead to clinical elimination. However, there are also *p*_1_*, p*_2_ combinations that lead to substantial tumor reduction on Day 25, but complete relapse by Day 150, e.g. *p*_1_ = 0.85*, p*_2_ = 0.36. It is also noteworthy that our Day 25 prediction that when *p*_1_ *> p*_2_, checkpoint blockade therapy will increase the proportion of low antigen tumor cells still holds on Day 150.

### 3.4 Clinical implications of variable antigen-mediated CTL proliferation rates

Figure 5 shows the changes in total tumor volume and composition on Day 25 and Day 150 for varying *α_nt_, α_mt_* combinations when the PD-1/PD-L1 checkpoint is active or completely blocked. Again, *α_nt_*is the maximum additional rate of CTL proliferation mediated by the presence of high antigen tumor cells and *α_mt_* is the maximum additional rate of CTL proliferation mediated by the presence of low antigen tumor cells.

**Figure 5:**
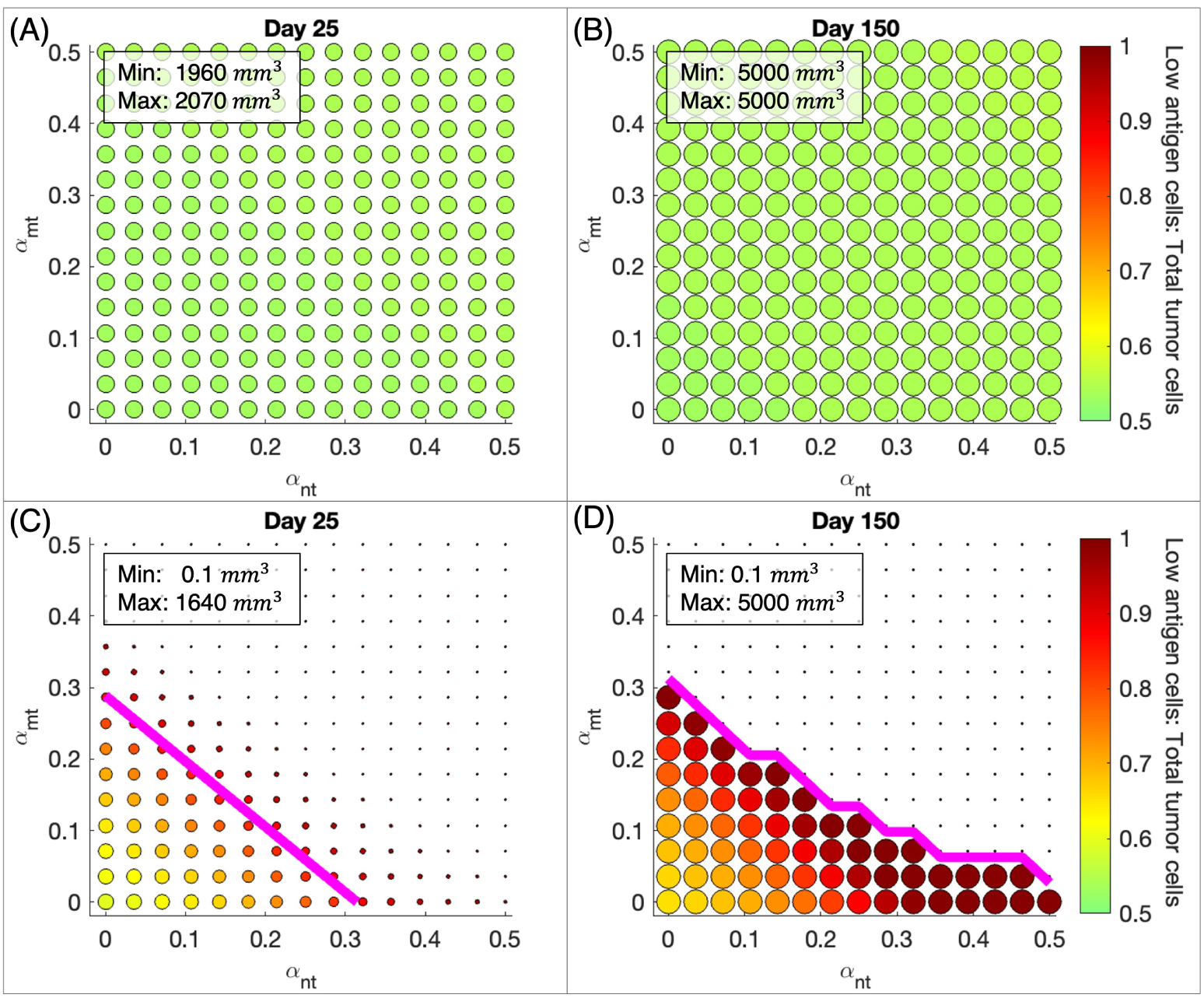
Combining ICI with therapy that modulates CTL proliferation activated by tumor cells is more effective in eliminating tumor than monotherapy. (A), (B), (C), (D) See Figure 4.

From Figure 5A, as *α_nt_, α_mt_* vary when the checkpoint is active, the total tumor volume on Day 25 remains largely unaffected, which is similar to what is shown in Figure 4A. In this cases, all tumors are within about 100 mm^3^ of each other, suggesting that changes in *α_nt_, α_mt_* cause slightly less variation in tumor size than *p*_1_*, p*_2_ on Day 25 when the checkpoint is active. By Day 150, all tumors have again reached the maximum carrying capacity volume (Figure 5B) when the checkpoint is active. In Figure 5, *p*_1_ and *p*_2_ are fixed at baseline values (0.92 and 0.33 respectively), which means that high antigen tumor cells have higher potential to perish via fast killing than low antigen tumor cells. Therefore, for all *α_nt_* and *α_mt_* values in all panels of Figure 5, the tumor shifts toward low antigen dominance on Day 25 and Day 150. This is in contrast to Figure 4 where the tumor can shift toward high or low antigen dominance depending on the *p*_1_*, p*_2_ values. When the checkpoint is active, Figures 5A and 5B show that tumor composition on Day 25 and Day 150 is largely similar despite variations in *α_nt_, α_mt_*, with the ratio of low antigen tumor cells to total tumor cells varying between 0.52 and 0.55.

Outcomes for complete checkpoint blockade are shown in Figures 5C and 5D. If tumors are characterized by low rates of T cell stimulation by tumor cells, the resulting tumor on Day 25 can grow to near 1/3 of the carrying capacity. However, tumors can be clinically eliminated when CTLs have a high rate of antigen-mediated proliferation, as illustrated by the difference in size of the dots in the upper right and lower left corners of Figure 5C. Tumors in Figure 5C are generally smaller than those in Figure 4C, suggesting that *α_nt_, α_mt_* may have a greater impact on final tumor size than *p*_1_*, p*_2_ on Day 25 when the checkpoint is blocked. Moreover, the proportion of low antigen tumor cells in the tumor increases significantly, as illustrated by the shift to more red colors in the in Figure 5C compared to Figure 5A. As *α_nt_, α_mt_* increase further, this shift toward low antigen dominance becomes more striking. Again, the magenta curve in Figure 5C represents the combinations of (*α_nt_, α_mt_*) that result in 75% reduction in total tumor volume on Day 25 and our baseline values fall into this region. If high antigen tumor cells stimulate T cell proliferation at our baseline rate of 0.15 per day, then low antigen tumor cells must stimulate T cell proliferation at a rate of at least 0.15 per day in order to achieve at least 75% tumor reduction on Day 25.

By Day 150 (Figure 5D),there is a complete split between two outcomes: escape or clinical elimination for all *α_nt_, α_mt_* combinations. There are also *α_nt_, α_mt_* combinations that lead to substantial tumor reduction on day 25, but complete relapse by Day 150, e.g. *α_nt_* = 0.21*, α_mt_* = 0.11. Similar to what Figure 3 suggests, increasing *α_nt_* and *α_mt_*leads to better therapeutic outcomes, with the other variables remaining the same. It is also noteworthy, that for all tumors that escape on Day 150 under checkpoint blockade therapy, there is an increased proportion of low antigen tumor cells as we see in Figure 4D.

### 3.5 Outcomes for virtual tumors before and after checkpoint blockade

To further investigate the effect of PD-1/PD-L1 checkpoint blockade on tumor reduction, we generate a virtual cohort of mice upon which to test therapeutic impact. Specifically, to predict the response of a diverse population comprising individuals with heterogeneous tumor and immune characteristics to checkpoint therapy, we use Latin Hypercube Sampling (LHS) to generate 30,000 combinations of all parameters, initial tumor composition and initial immune to tumor cell ratio. Each parameter combination represents a mouse with unique tumor and immune dynamics. We categorize our simulated outcomes before and after complete checkpoint blockade into clinical elimination (tumor size less than 0.1mm^3^), dormancy (tumor size between 0.1mm^3^ and 500 mm^3^) and escape (greater than 500 mm^3^). Figure 6A shows that when the checkpoint is active approximately 20% of tumors are small enough to be dormant on Day 25, but almost 80% are already large enough to be in the escape category. By Day 150, almost all dormant tumors have progressed to escape with only a very small fraction being clinically eliminated as shown in Figure 6B.

**Figure 6:**
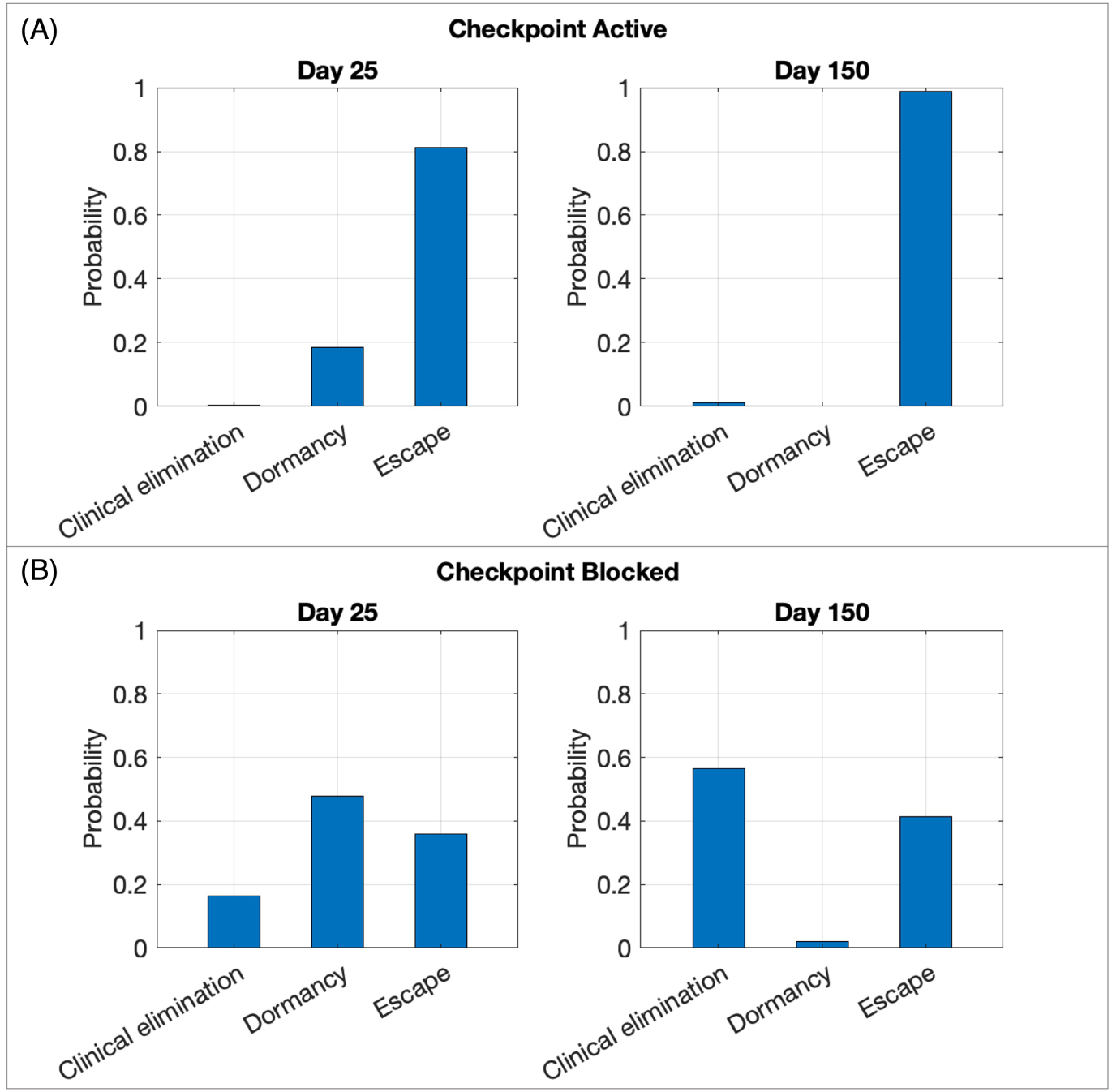
Virtual cohort simulations show that checkpoint blockade results in better clinical outcomes on both Day 25 and Day 150. (A) Probability of each clinical outcome when the checkpoint is active. (B) Probability of each clinical outcome when the checkpoint is blocked. Clinical elimination: tumor size *<* 0.1 mm^3^; dormancy: 0.1 *≤* tumor size *≤* 500 mm^3^, escape: *>* 500 mm^3^). Size of virtual cohort: 30,000.

When the checkpoint is completely blocked, Figure 6C shows that on Day 25 about 16% of tumors are clinically eliminated and only about 36% have escaped. By Day 150, most tumors that were dormant on Day 25 have been clinically eliminated and a small amount have escaped, as shown in Figure 6D. Taken together, the results illustrated in Figure 6 suggest that Day 25 tumor size alone can be a poor predictor of long-term clinical outcomes, especially when the tumor is small but detectable.

### 3.6 Correlation between key immune parameters and therapeutic outcomes

We showed in the previous section that mice with different tumor-immune landscapes have distinct responses to checkpoint therapy. In Figure 7 we investigate how clinical outcomes depend on the values of important parameters, which relate to specific tumor and immune characteristics in our cohort of 30,000 mice. Figures 7A, C, E, and G are stacked bar plots showing the probability of each outcome (clinical elimination, dormancy, or escape) for a range of a chosen parameter that impacts clinical outcome. Figures 7B, C, F, and H are violin plots showing the distribution, median and interquartile range of parameters in our virtual cohort associated with each outcome.

**Figure 7:**
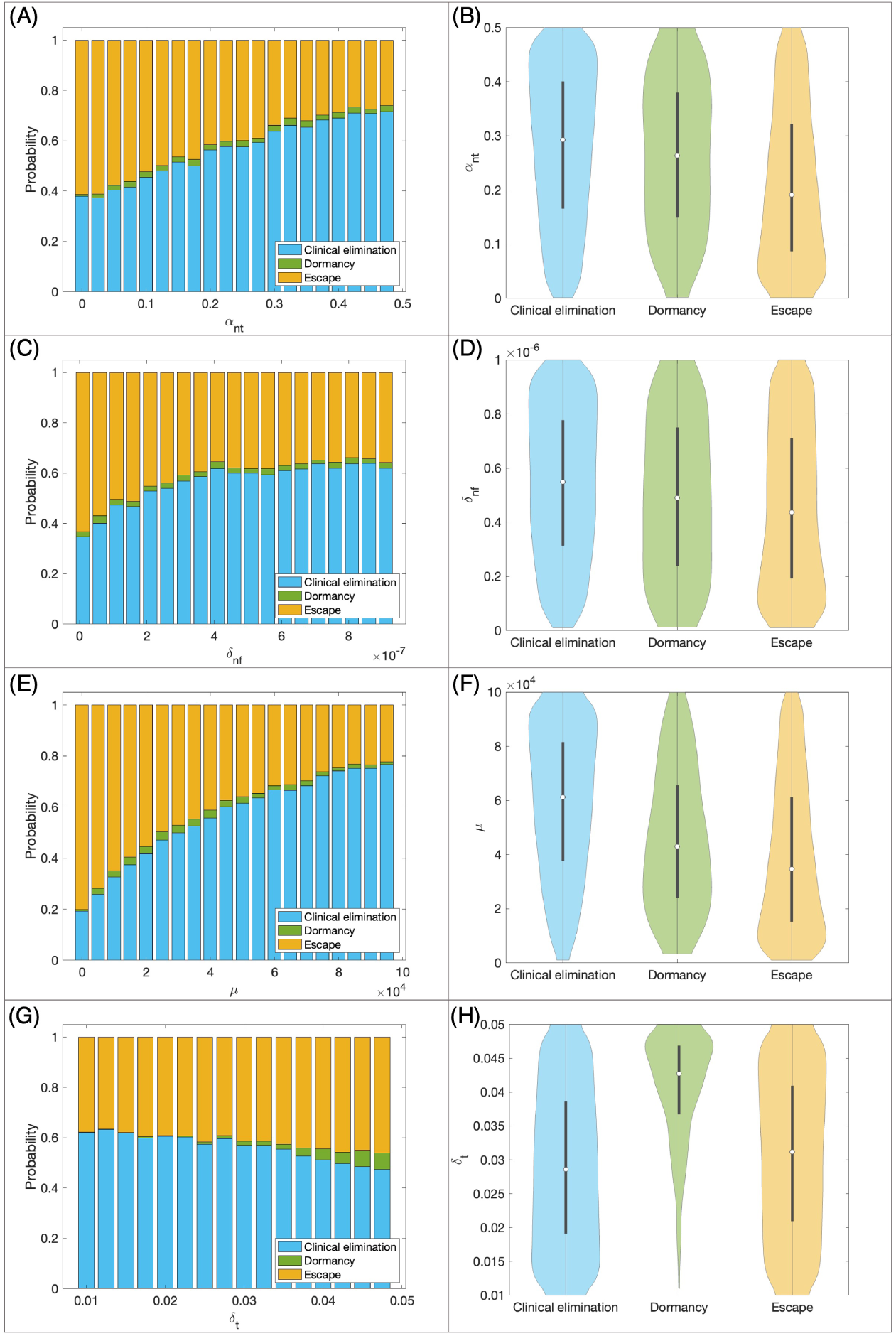
Relationship between model parameters and the outcome of checkpoint blockade therapy in the same virtual cohort as Figure 6. (A),(C),(E),(G) Stacked bar plots of the probability of each outcome (clinical elimination, dormancy, or escape) for a range of a chosen impactful parameter. (B),(D),(F),(H) violin plots of the distribution of parameters of the virtual cohort associated with each outcome, with the shape showing probability density, the white circle showing median and the grey lines showing interquartile range.

Figure 7A implies that as the maximum rate of CTL proliferation activated by high antigen tumor cells (*α_nt_*) increases from smallest values to the largest, the probability of clinical elimination also increases from 0.38 to 0.72, while the probability of escape decreases from 0.61 to 0.26. In our cohort, the *α_nt_* median and interquartile range for escape cases are lower than that of the clinical elimination and dormancy cases as shown in Figure 7B. CTL-induced death rate of high antigen tumor cells via the fast-killing mechanism (*δ_nf_*) and activation and recruitment rate of T cells (*µ*) show similar trends (Figures 7C,D,E,F). However, the trend is more moderate for *δ_nf_* but more pronounced for *µ*. As *µ* increases from the smallest to the largest values, the probability of elimination quadruples while the probability of elimination drops down to one fourth. Moreover, the *µ* median and interquartile range of elimination cases is also significantly higher than that of escape cases. These all reiterate the importance of *µ* in tumor elimination as we explained in 3.2.

Regarding the death rate of T cells (*δ_t_*), we observe that the probability of clinical elimination decreases slightly and probability of escape increases slightly as *δ_t_* increases (Figure 7G). We also see that the median and interquartile range for *δ_t_* in our virtual cohort for dormancy cases is higher than that of clinical elimination and escape cases, which means that dormancy is extremely unlikely with small *δ_t_* and its probability improves slightly with bigger *δ_t_* values.

## 4 Discussion

Here we presented the first mechanistic mathematical model designed to investigate the impact on immunotherapy outcomes of differential cell-kill strategies immune cells use to target tumor cells with different antigen loads. To predict which patients are more likely to respond to ICIs, and to develop improved therapeutic strategies, understanding the differential cell-kill mechanisms that T cells use against tumor cells is essential. Since the discovery of the PD-1/PD-L1 immune checkpoint, the immunosuppressive effect of the checkpoint has been included in several mathematical models of tumor-immune dynamics [20, 22, 31, 23]. While our model has similar features to existing models, our work is distinctive and innovative in two ways: i) we consider two types of tumor cells (high antigen and low antigen phenotype), and ii) we explicitly incorporate the two killing mechanisms.

We assessed the impact of ICI monotherapy or combination therapy by analyzing the relationship of important model parameters to tumor volume and composition. Based on bifurcation analysis of potential long-term tumor responses, we categorized therapeutic outcomes into “elimination”, “dormancy” and “escape”, which can correspond to the three phases of the immunoediting framework.[24]. “Elimination” referred to a tumor volume of 0 mm^3^, while “dormancy” referred to a tumor with volume between 0.1 and 500mm^3^ and “escape” refers to a tumor expanding larger than 500mm^3^. To better understand how parameters impact therapy outcomes in a more clinically relevant time frame, we supplemented bifurcation analysis with direct visualization of Day 25 and Day 150 tumor volume and composition with varying parameters in Figure 4 and 5. We chose the endpoints, Day 25 and Day 150, to reflect immediate and long-term responses to therapy, respectively. In this case, we re-coined “elimination” as “clinical elimination” to refer to a clinically undetectable tumor with volume less than 0.1mm^3^, representing the limit of clinical and imaging assessment.

A novel feature of our model was that the probability of tumor cell death via fast killing differs based on tumor antigenicity, with *p*_1_ being the probability associated with high antigen tumor cells and *p*_2_ associated with low antigen tumor cells. Our analysis showed that, after checkpoint blockade therapy, *p*_1_ and *p*_2_ critically determine whether tumor elimination is possible and how tumor composition changes if the tumor persists. In Figure 4, we showed that if the probability of tumor cell death via fast killing is sufficiently high for both high and low antigen tumor cells, ICI monotherapy can clinically eliminate the tumor. Under our baseline assumption, high antigen tumor cells are more likely to be killed via the fast mechanism than low antigen tumor cells, i.e. *p*_1_ *> p*_2_. In this case, when the tumor persists under ICI monotherapy, the resulting tumor is more low antigen-dominant than the initial tumor in the short term. The tumor then progresses to become wholly composed of low antigen tumor cells over time. If our baseline assumption holds, our result suggested that the presence of low antigen tumor cells weakens the response to ICI monotherapy. A related phenomenon was observed clinically in a recent study by Zapata et al. [36], which demonstrated that immune-edited tumors with low tumor antigenicity are less likely to respond to ICIs.

Furthermore, we studied how to enhance the therapeutic results of ICI with combination therapy. To do this, we focused on parameters that can be altered by current clinical interventions. CTL activation and recruitment rate (*µ*) and maximum rate of antigen-mediated CTL proliferation (*α_nt_, α_mt_*) proved to be the most important immune parameters for achieving tumor reduction or elimination. We showed that increasing *µ* can lead to a much smaller tumor or even elimination, with other tumor-immune characteristics kept constant. One way to increase CTL activation and recruitment is through adoptive T-cell therapy. Emerging clinical evidence supports the use of adoptive cell therapy using tumor-infiltrating lymphocytes after anti-PD-1 or PD-L1 therapy [27, 7], as well as the use of CAR T-cell therapy before anti-PD-1 to treat solid tumors [1]. Moreover, according to our simulations, increasing *α_nt_* or *α_mt_* can also lead to significant volume reduction. IL-2, a cytokine that promotes the growth and activation of T cells, is often used in combination with other forms of immunotherapy, including adoptive T-cell transfer [17]. Therefore, cytokine therapy can be potentially combined with ICIs to increase antigen-mediated T cell proliferation (*α_nt_*, *α_mt_*) and produce better therapeutic outcomes. Using a humanized mouse model, Jespersen et al. determined that continuous presence of IL-2 is essential for eradicating tumors undergoing adoptive T-cell transfer and/or anti-PD1 [16], although they did not manage to show that anti-PD-1 and adoptive T-cell transfer combination therapy is superior than adoptive T-cell transfer alone. In practice, IL-2 therapy combined with ICI has not proven more effective than ICI monotherapy to date [32], although a recent phase 1b study suggested that combination therapy with high-dose IL-2 therapy and anti-PD-1 might be feasible and tolerable [30]. Nonetheless, new variants of IL-2 for synergy with ICI continued to be developed in clinical setting [32]. Overall, combination therapy may have two significant benefits: i) achieve tumor elimination or drastic tumor volume reduction that is otherwise unattainable with ICI alone; ii) reduce the amount of drug used in each treatment, which patients might tolerate better.

Beyond immunotherapy alone, our analysis showed that ICIs can also be combined with firstline treatments like chemotherapy to produce therapeutic benefits. The “elimination or escape” and “dormancy or elimination” bistability regions in Figure 3 suggested that, depending on the initial tumor size, tumor composition, or ratio of tumor to immune cells, two patients with similar tumor-immune landscapes may have tumors of vastly different sizes after receiving checkpoint blockade therapy. Furthermore, mouse models showed that chemotherapies that induce immunogenic cell death can turn a non-T cell-inflamed tumor into a T cell-inflamed tumor more infiltrated with tumor-specific T cells [10]. Some chemotherapies can upregulate the expression of PD-L1 by cancer cells [10]. Since T cell-inflamed tumors elicit different immune responses from non-T cell-inflamed tumors, which would translate to different immune or tumor parameters in our model, ICI can produce qualitatively different therapeutic outcomes depending on the baseline endogenous immune response to the tumor. Therefore, chemotherapy may be used before ICI to enhance the efficacy of ICI and facilitate tumor elimination. This strategy has been successful clinically bladder cancer where maintenance therapy with avelumab, an anti-PD-L1 therapy, has become standard of care with an overall survival benefit [25].

To single out the impact of each aforementioned model parameter on an individual patient, we had to keep the other parameters invariant. Through virtual cohort generation and simulations, we shifted our focus to a diverse population and study the relationship between specific characteristics of the tumor-immune landscape and the outcomes of checkpoint blockade therapy or combination therapy. High CTL recruitment rate (*µ*), antigen-mediated T cell proliferation rates (*α_nt_, α_mt_*), and CTL-induced death rate of tumor cells via fast killing (*δ_nf_, δ_mf_*) all corresponded to a higher probability of tumor elimination after checkpoint blockade therapy. In particular, this was most pronounced for *µ*: patients with a high CTL recruitment rate (*µ*) were four times more likely to their tumors eliminated than those with a low *µ*. If these patient-specific parameters can be measured, they can be used to predict the probability of tumor elimination after receiving ICI alone. If other forms of cancer treatments (e.g. adoptive T-cell transfer, cytokine therapy) can modulate these parameters, they should be considered for combination therapy with ICI for better odds of tumor elimination. The correlation between the rate of fast killing for high and low antigen tumor cells (*δ_nf_, δ_mf_*) and the outcomes of checkpoint blockade therapy highlighted the importance of considering differential immune cell-kill mechanisms when evaluating the efficacy of ICI. Furthermore, virtual cohort simulations suggested that it might be too soon to conclude the outcome of ICI on Day 25, because tumors that shrunk significantly by Day 25 might relapse by Day 150, as implied by results in Figures 4, 5, and 6.

This study thoroughly investigated the outcomes of complete immune checkpoint blockade as the best-case scenario for using ICIs. Implementing a realistic dosing schedule of anti-PD-1 therapy in the virtual cohort will improve the clinical relevance of our model. Despite the computational and analytical advantages of continuous ODEs in modeling tumor-immune dynamics, the lack of spatial components in ODEs lead to the inability to obtain structural information about the tumor and the tumor microenvironment. Therefore, agent-based models, which describe each cell as an independent agent in three-dimensional space and prescribe how cells move or interact, can complement ODE models. These approaches, along with parameterization of ODE models with in vivo data in the future, will improve representation of divers patient populations in the models.

Data-driven and biologically informed mathematical models of cancer control strategies are a powerful complement to experimental studies. Mechanistic ODE models like the ones presented in this work allow rapid simulations to identify critical patterns or discover underlying mechanisms in the tumor microenvironment that drive cancer progression and therapeutic resistance. In particular, we explored how differential cell-kill mechanisms that T cells use against tumor cells with variable antigenicity impact tumor growth and their implications on ICI monotherapy or combination therapy. Our methodology can be used to systematically explore a wide range of questions related to tumor-immune dynamics and immunotherapy. The modeling framework and extensions proposed can provide valuable insights for the rational design of pre-clinical experiments and clinical trials.

