## Supplemental Figure 1 for "Mathematical Model Predicts Tumor Control Patterns Induced by Fast and Slow CTL Killing Mechanisms"

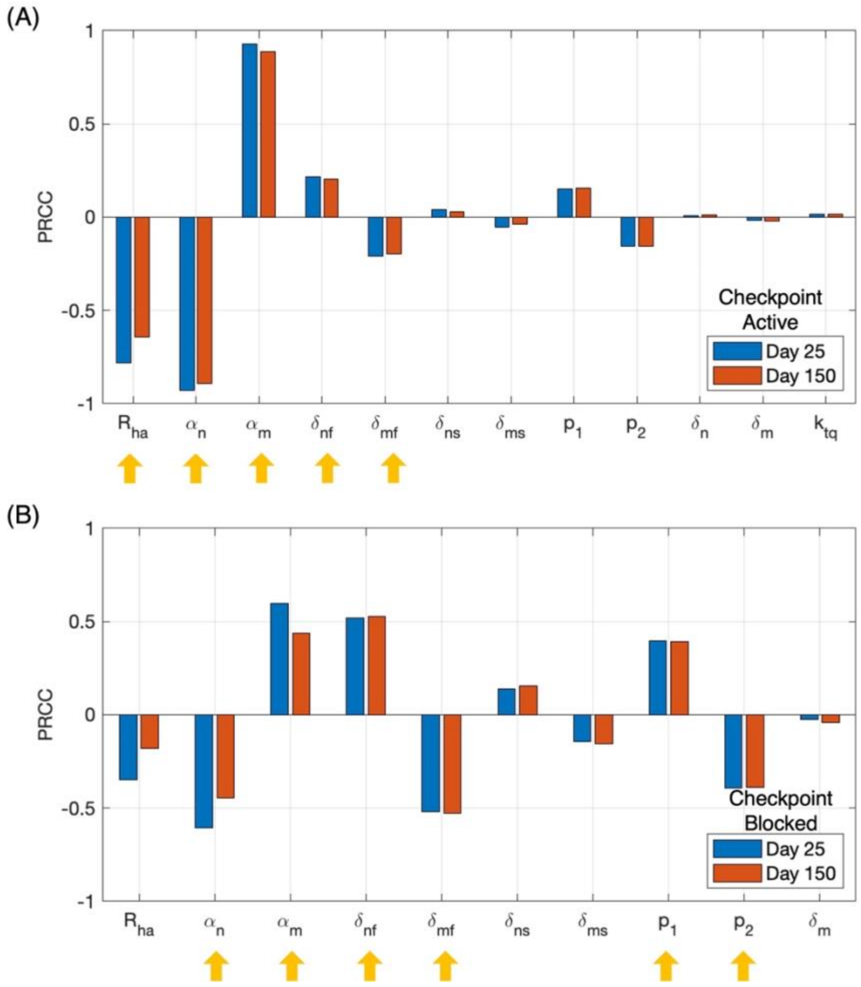


Figure S1: Immune checkpoint activity affects the most sensitive parameters with respect to ratio of low antigen tumor cells to total tumor cells. (A) PRCCs (partial rank correlation coefficient) of parameters in the model with immune checkpoint active. (B) PRCCs of parameters in the model with immune checkpoint blocked. Blue: PRCC with respect to the ratio of low antigen tumor cells to total tumor cells on Day 25. Red: PRCC with respect to the ratio of low antigen tumor cells to total tumor cells on Day 150. Yellow arrows: sensitive parameters with magnitude of PRCC ranked in the top quartile. Parameters with magnitude of PRCC ranked in the top 50% are shown.
